# Heterotic Pooling Strategy for Enhanced Hybrid Performance in Bitter Gourd (*Momordica charantia* L.)

**DOI:** 10.1101/2025.09.24.678193

**Authors:** Anita Verma, Shikha Dubey, Alpana Joshi

## Abstract

Bitter gourd (*Momordica charantia* L.), a crop of significant nutritional and medicinal renown, is experiencing a surge in global demand, particularly in Asia. To meet this demand amidst the challenge of limited agricultural land, enhancing productivity through innovative breeding is essential. Hybrid breeding, which harnesses the power of heterosis, offers a proven strategy for boosting crop yields. However, the success of any heterosis breeding program, especially in a naturally cross-pollinated and genetically variable crop like bitter gourd, is fundamentally dependent on robust methods for assessing and strategically utilizing the available genetic diversity. This study compared the efficacy of traditional morphological trait-based clustering with a modern breeding value-based method (Heterotic grouping based on Specific and General Combining Ability - HSGCA) for predicting heterosis. Eleven diverse bitter gourd inbred lines were crossed in a half-diallel fashion to generate 55 F1 hybrids. The hybrids and parents were evaluated for key yield-related traits. K-means clustering based on morphological data resulted in five groups but proved unreliable for predicting heterosis, as high heterosis was often observed within clusters. In contrast, HSGCA-based clustering organized the lines into three coherent groups, where crosses between the two main groups (Group 1 and Group 2) consistently exhibited significantly higher mid-parent (20.89%) and better-parent (7.5%) heterosis compared to intra-group crosses. The HSGCA method provides a far more reliable framework for defining heterotic groups in bitter gourd, enabling more efficient and predictive hybrid breeding programs.

## 1. Introduction

Bitter gourd (*Momordica charantia* L.), a crop of significant nutritional and medicinal renown, is experiencing a surge in global demand, particularly in Asia. To meet this demand amidst the challenge of limited agricultural land, enhancing productivity through innovative breeding is essential. Hybrid breeding, which harnesses the power of heterosis, offers a proven strategy for boosting crop yields. However, the success of any heterosis breeding program, especially in a naturally cross-pollinated and genetically variable crop like bitter gourd, is fundamentally dependent on robust methods for assessing and strategically utilizing the available genetic diversity.

Over the years, researchers have employed a suite of statistical tools to characterize bitter gourd germplasm. Phenotypic diversity has been extensively quantified using Mahalanobis D^2^ analysis (Tyagi et al., 2017; Maurya et al., 2018; Bhati et al., 2023; Verma & Joshi, 2025) and Principal Component Analysis (PCA) (Mallikarjun et al., 2023) to group genotypes. More recently, molecular techniques, especially the use of SSR markers, have provided higher-resolution insights, revealing genetic clusters often linked to geographic origin and key fruit traits (Saxena et al., 2015; Alhariri et al., 2021; Meghashree et al., 2024). While these methods are invaluable for assessing genetic distance, their utility in directly predicting hybrid performance is fraught with uncertainty.

A critical limitation, as highlighted in numerous studies, is the inconsistent and often poor correlation between genetic distance and heterosis. The underlying reasons are complex; as Yu et al. (2005) noted, not all divergent loci contribute equally—or at all—to heterosis. Furthermore, complex traits like yield are often governed by a few major QTLs, meaning that overall genetic distance is not a reliable proxy for the divergence at these specific, crucial loci responsible for yield gain (Yuan et al., 2008; Pan et al., 2020). A comprehensive review by Dias et al. (2004) confirmed this inconsistency across 54 studies, showing the relationship between genetic distance and hybrid performance to be variable. Given the inherent limitations of predicting hybrid performance from simple genetic distance, the need for more innovative and reliable approaches to group germplasm has become increasingly apparent. To overcome the constraints of conventional methods, which often rely on morphological diversity followed by extensive validation, researchers have shifted focus towards utilizing breeding values—specifically General Combining Ability (GCA) and Specific Combining Ability (SCA)—to classify parental lines into effective heterotic groups. This shift acknowledges that heterosis arises from a complex interplay of both additive (GCA) and non-additive (SCA, dominance, epistasis) genetic effects. As studies have shown, many high-performing hybrids result from crosses where at least one parent has low GCA, underscoring the importance of specific combinations (Liu et al., 2022).

A key innovation in this area is the Heterotic grouping based on Specific and General combining ability method proposed by Fan et al. (2009), which integrates both GCA and SCA to more accurately predict a line’s potential to produce superior hybrids. This approach has demonstrated significantly higher breeding efficiency compared to traditional methods. To address the limitation of focusing on a single trait like yield, the HGCAMT method was later proposed, incorporating GCA effects from multiple traits to provide a more comprehensive assessment, especially under stress conditions (Badu-Apraku et al., 2013).

Subsequent comparative studies have consistently highlighted the efficacy of these breeding value-based methods. Adewale et al. (2023) found HSGCA to be the most accurate (87% success rate) and highly efficient (52.8% breeding efficiency) method for classifying maize lines, performing comparably to high-throughput DArTseq markers. This superiority of HSGCA is supported by research in various crops, including baby corn (Kumar et al., 2022) and sweet corn (Mahato et al., 2021). However, the literature also indicates that the most effective method can be context-dependent, varying with the specific germplasm and environment (Badu-Apraku et al., 2015, Badu-Apraku et al., 2016; Oyetunde et al., 2020). This underscores the critical need for empirical validation within a specific crop and breeding program.

In light of these advancements, the present study was undertaken in bitter gourd to address this need by conducting a direct comparative analysis to determine whether a modern, breeding value-based approach (HSGCA) offers a more reliable framework for predicting heterosis than traditional clustering based on morphological traits.

## 2. Material and Method

### 2.1. Germplasm

This study began with 22 distinct open-pollinated (OPV) bitter gourd varieties sourced from across India. A significant number of these came from West Bengal and Odisha, a region known as a primary center for this crop’s diversity. After analysis for Distinctness, Uniformity, and Stability (DUS), 11 lines with consistent and stable physical traits were selected for the next phases of the research (Table 1).

**Table 1.** List of Material used in the heterosis and combining ability study.

| S. N. | Name | Code | Collected Form |
| --- | --- | --- | --- |
| 1 | KAJAL | P1 | Odisha |
| 2 | KARISHMA | P2 | Odisha |
| 3 | KUHELI | P3 | Odisha |
| 4 | NILIMA | P4 | Odisha |
| 5 | TIYASHA | P5 | Odisha |
| 6 | BG-0008 | P6 | Maharashtra |
| 7 | BG-0002 | P7 | Maharashtra |
| 8 | KATAHI | P8 | UP |
| 9 | BOLDERUCCHA | P9 | West Bengal |
| 10 | KALOSONA | P10 | West Bengal |
| 11 | RSMEGHANA2 | P11 | West Bengal |

### 2.2. Generation of Diallel Crosses

Using a half-diallel mating design, the 11 selected bitter gourd parental lines were manually crossed to generate 55 F1 hybrids. The process began the evening before flowering when designated male and female flower buds were enclosed in butter paper bags to prevent unwanted pollination. The following morning, between 6:00 and 8:00 AM, pollen was transferred from the male parent to the female parent’s stigma. After this hand-pollination, the flowers were labelled and re-bagged. Finally, seeds were extracted from the resulting fruits and saved for growing the F1 generation in the subsequent season.

### 2.3. Experimental Location and Design

The study was conducted during the Kharif season of 2022 at the BRDC Centre of Bayer Crop Science, located in Kallinayakanhalli, Bangalore, Karnataka, India. Eleven diverse bitter gourd parental lines were used to generate 55 F1 hybrids in a half-diallel fashion, excluding reciprocals. This crossing design allowed for the evaluation of both general combining ability (GCA) and specific combining ability (SCA) of the parental lines. The experimental design was a Randomized Complete Block Design (RCBD) with three replications. This design allowed for the control of environmental variation and ensured that the results were statistically sound.

### Data Studied for Heterosis and Combining ability

To conduct the heterosis and combining ability analyses, 55 generated hybrids, along with 11 parental lines, were evaluated in three replications. Data were collected for five key traits related to fruit quality and yield. These traits were chosen based on their economic importance and relevance to consumer preferences.

#### Average Fruit Weight (AFW) (g)

For each genotype, 10 fruits were randomly selected from the third harvest, and their weight was measured using an electronic balance. The average weight of these fruits was then calculated.

#### Early Yield (EYDPA) (tons/acre)

Early yield was defined as the cumulative weight of mature fruits harvested from the first two pickings. This metric reflects the yield obtained when produce typically commands a higher market price.

#### Total Yield (YDPA) (tons/acre)

Total yield was calculated by summing the weight of all mature fruits harvested throughout the entire picking period. A total of ten pickings were conducted, and the crop was maintained for 110 days.

#### Fruit Length (FRLGT) (cm)

The length of 10 randomly selected red-ripe fruits from the third harvest was measured in centimetres, from the base to the tip of each fruit. The average of these measurements was recorded as the fruit length.

#### Fruit Girth (FRGTH) (cm)

Fruit girth was measured at the midpoint of the same 10 fruits used for fruit length measurements, using a Vernier calliper. The average girth was then calculated.

### 2.6. Statistical analysis

#### Combining Ability Analysis

GCA and SCA effects and their variances were determined as per Griffing method-II (Griffing, 1956a). This method includes F_1_ (all possible combinations between parental lines made in half diallel fashion) and parents themselves.

From the (E.M.S.) given in Table 2, the estimation of variance components of GCA (σ^2^gca) and SCA (σ^2^sca) were calculated as follows:

**Table 2:** Outline of ANOVA and Expected Mean Squares (Griffing’s Method-II)

| Source of Variation | Degrees of Freedom (df) | Mean Square (MS) | Expected Mean Square (EMS) |
| --- | --- | --- | --- |
| GCA | $p - 1$ | MSG | $\sigma^2e + 2\sigma^2sca + (p+2)\sigma^2gca$ |
| SCA | $p(p - 1) / 2$ | MSS | $\sigma^2e + 2\sigma^2sca$ |
| Error | $(r-1)(p(p+1)/2 - 1)$ | MSE | $\sigma^2e$ |

NOTE: p is the number of parents, r is number of replications, MSG: Mean Square for GCA, MSS: Mean Square for SCA, MSE: Mean Square for Error.

GCA Variance (σ^2^gca): σ^2^gca= (MSG - MSS) / (p + 2)

SCA Variance (σ^2^sca): σ^2^sca = (MSS - MSE) / 2

Estimation of GCA effects, g_i_ was calculated using below equation as per Griffing Method II:

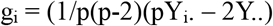

Estimation of SCA effects, s_ij_ was calculated using below equation as per Griffing Method II:

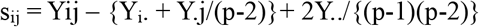

1. **Y**_**i**_.: This represents the *sum of all observations (crosses)* that involve parent *i*. Therefore, Yi. = Y_□□_ + Y_□□_ + Y_□□_ +… +Y_□□_
2. **Y.j:** This represents the *sum of all observations (crosses)* that involve parent *j*.
3. **Yij**: Mean of parent i cross with parent j
4. **Y**..: Represents the grand total of all observations
5. **p**: The number of parents.

#### HSGCA Value and Clustering

The inbred lines were assigned into heterotic groups based on the phenotypic diversity and heterotic group’s specific and general combining ability (HSGCA) for yield. The HSGCA was computed as follows: HSGCA= Cross mean Xij−Tester mean (Xi) = GCA+ SCA

Where Xij is the mean yield of the cross between ith tester and jth line, Xj is the mean yield of the ith tester and X.j is the mean yield of jth line.

### 2.7. Statistical Analysis for Diversity Assessment

#### K-Mean Clustering

To assess the genetic diversity and classify the 11 bitter gourd genotypes, K-means clustering was employed to partition the genotypes into a pre-defined number of distinct, non-overlapping clusters. This analysis was performed twice using different input data: first with standardized morphological trait data, and subsequently with the calculated General and Specific Combining Ability (HSGCA) values. The optimal number of clusters (K) for each dataset was determined using the elbow method, which identifies the point on a scree plot where the rate of variance explanation diminishes significantly with the addition of more clusters. The K-means algorithm itself iteratively assigned genotypes to clusters based on Euclidean distance to randomly initialized cluster centers, recalculating the centroids after each assignment until cluster membership stabilized. The final groupings for both analyses were visualized using scatter plots, where the genotypes were plotted along the first two principal components and color-coded by their assigned cluster.

#### Statistical analysis for validating heterotic groups

To validate the efficiency of different heterotic grouping methods, mid-parent heterosis between and within groups was utilized. Heterosis was then calculated as the percentage by which the F_1_ hybrid’s mean performance differed from the mean performance of parental lines used in the cross. The specific calculations for these heterosis percentages were based on the methods described by Turner (1953) and Hayes et al. (1955).

#### Mid-Parent (MP) Heterosis

This is the percentage difference between the mean performance of F_1_ hybrid and the *average* performance of its two parents (MP). A positive value indicates the hybrid outperformed the average of its parents.

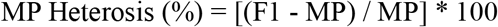

## 3. Results

A K-means clustering analysis was conducted on the morphological data to sort the 11 genotypes into a set number of discrete groups, providing a more conclusive classification.

### 3.1. Phenotypic Diversity and Trait-Based Clustering

Initial assessment of the germplasm focused on morphological traits. To objectively group the 11 genotypes based on their physical characteristics, a K-means clustering analysis was performed. The scree plot analysis (Fig. 1) indicated that the optimal number of distinct groups within the dataset was five (K=5). The resulting cluster plot (Fig. 2) visually represents the separation of the genotypes into these five groups based on the first two principal components.

**Fig 1:**
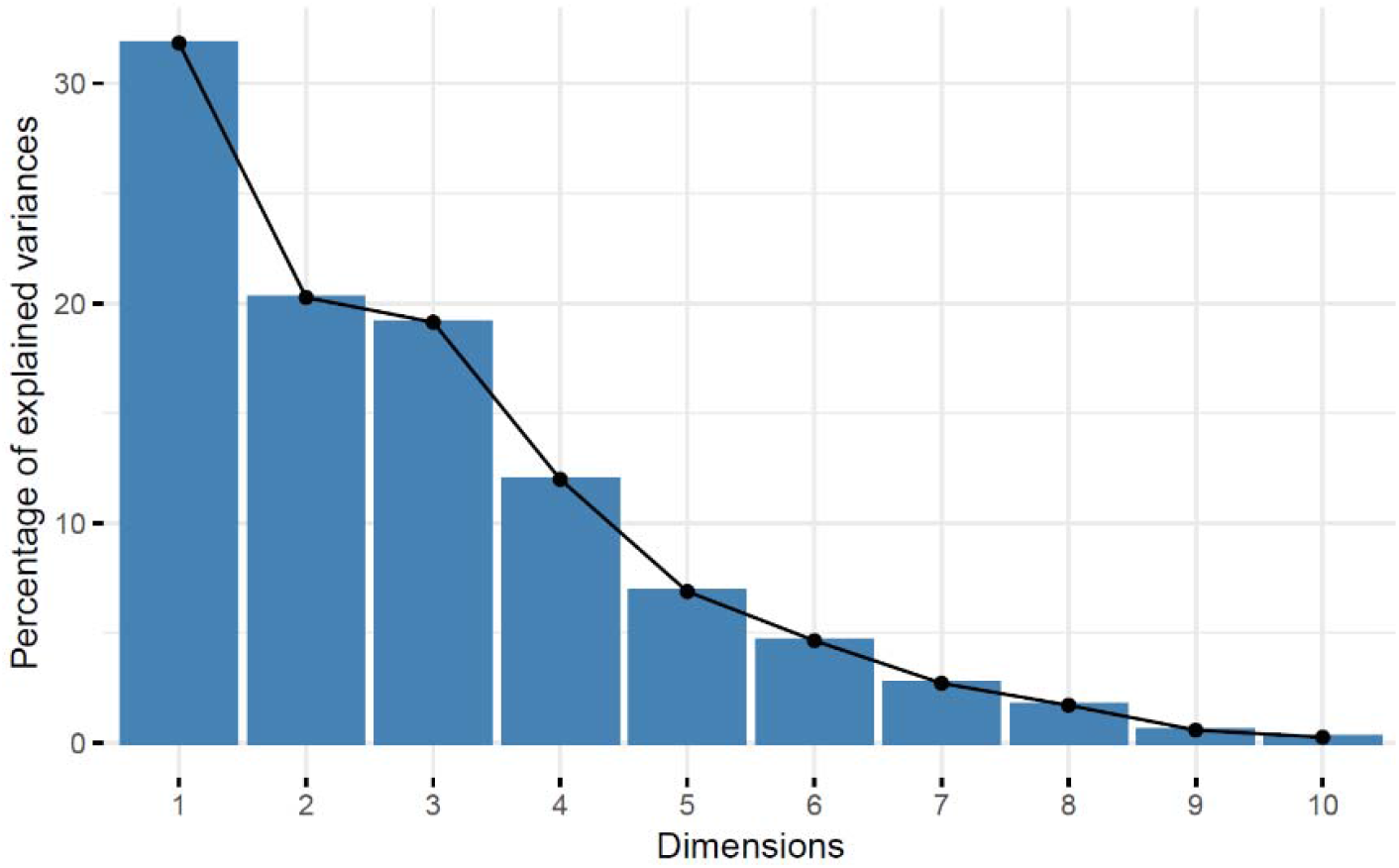
Scree Plot for Determining the Optimal Number of Groups.

**Fig 2:**
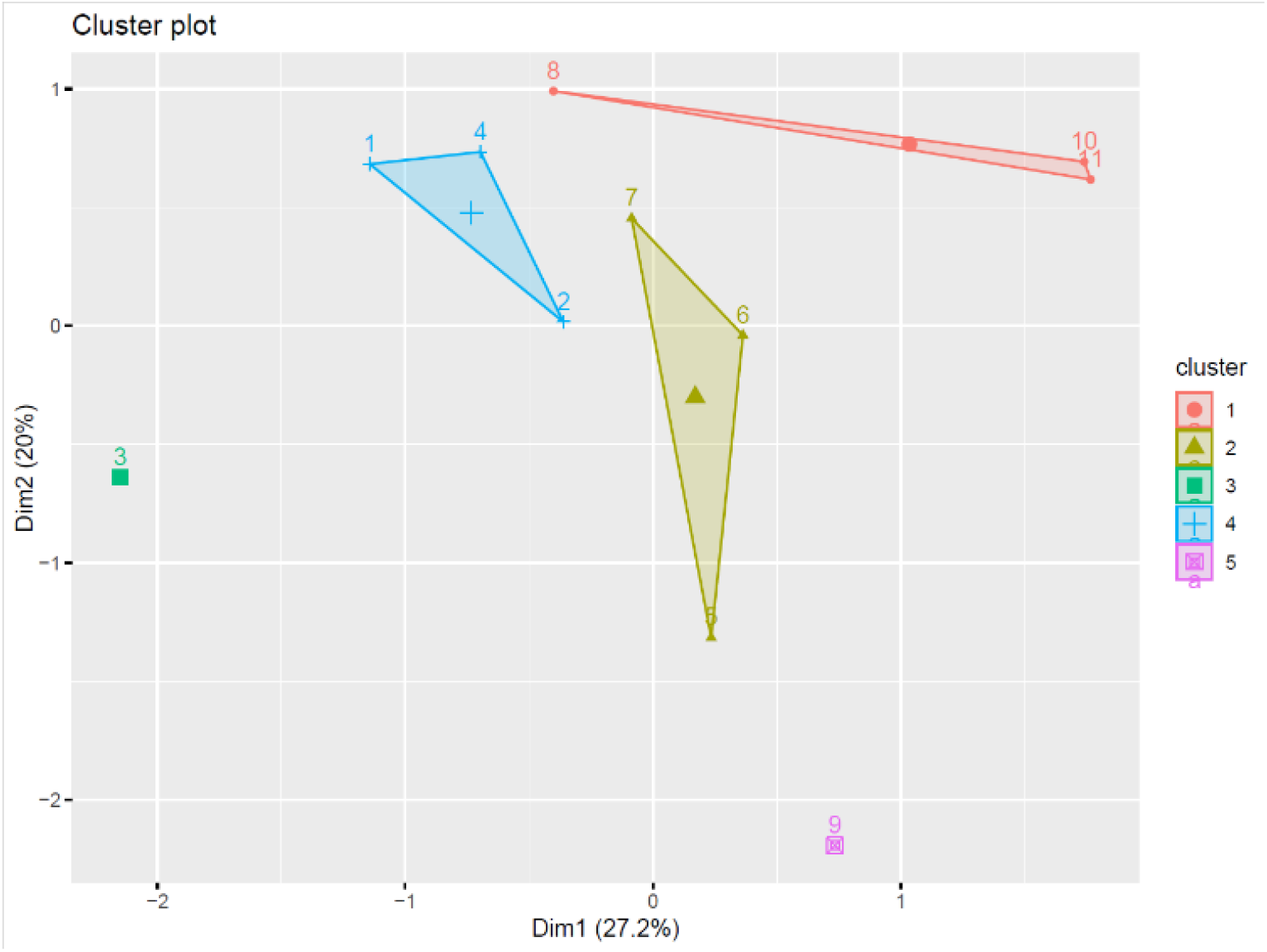
K Mean Clustering using phenotypic data.

The final composition of the five phenotypic clusters is detailed in Table 3. Group 1 contained three genotypes (P8, P10, P11), Group 2 also contained three (P5, P6, P7), and Group 4 contained three genotypes from Odisha (P1, P2, P4). Notably, two genotypes, P3 (KUHELI) and P9 (BOLDERUCCHA), were each segregated into their own distinct clusters (Group 3 and Group 5, respectively). This isolation confirmed their status as morphologically unique accessions within the panel.

**Table 3:**
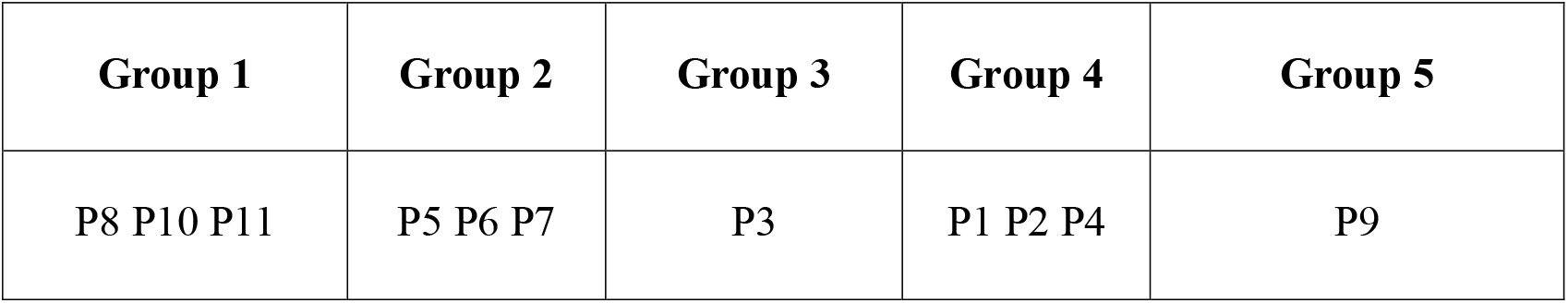
Distribution of genotypes into different clusters based on morphological traits using K-Means Analysis.

| Group 1 | Group 2 | Group 3 | Group 4 | Group 5 |
| --- | --- | --- | --- | --- |
| P8 P10 P11 | P5 P6 P7 | P3 | P1 P2 P4 | P9 |

### 3.2. Genetic Potential Revealed by Combining Ability Analysis

Recognizing that phenotypic similarity does not guarantee a corresponding breeding value, the focus of the investigation was transitioned. The ultimate utility of a genotype for a plant breeder lies in its ability to produce superior offspring, a quality best measured by its combining ability. Therefore, to assess the genotypes based on their genetic potential, a subsequent K-means analysis was conducted using GCA and SCA values (HSGCA) as input (Table 4). The core objective was to create a new set of clusters based on breeding performance and to conduct a comparative analysis against the morphology-based clusters. This comparison was designed to reveal which diversity assessment strategy is more powerful for predicting heterotic combinations.

**Table 4:**
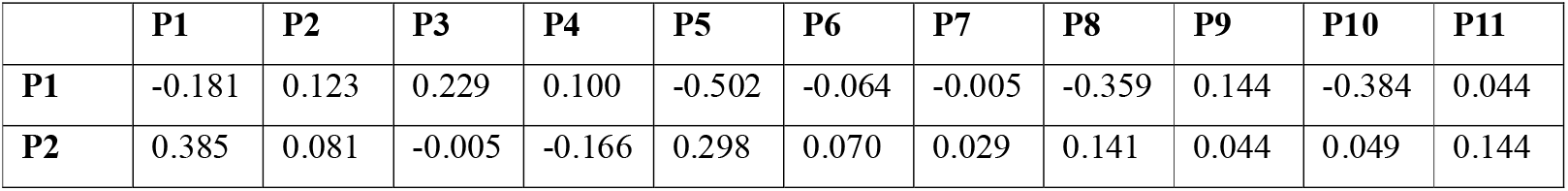
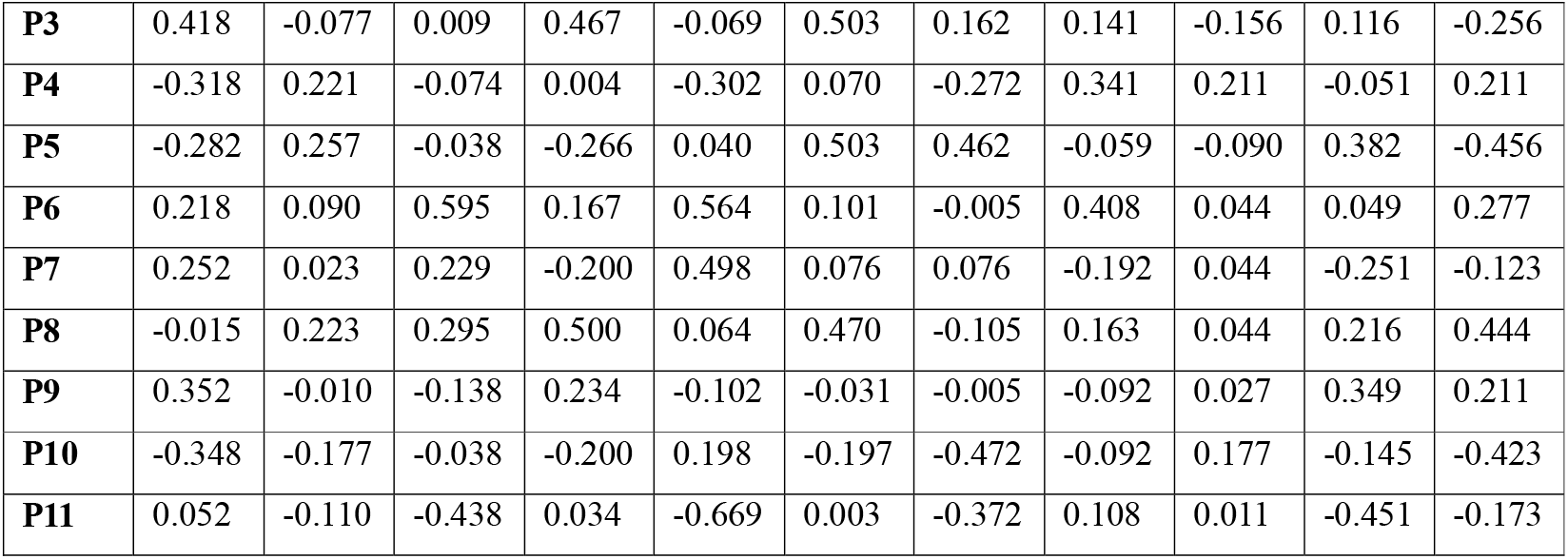
HSGCA Value (GCA+SCA) for all pairwise crosses.

The application of K-means clustering to the combining ability data resulted in the classification of the 11 genotypes into three distinct groups, providing a clear framework based on breeding value. The composition included two multi-genotype clusters—Group 1 (P1, P4, P10, P11) and the largest, Group 2 (P2, P3, P6, P7, P8, P9)—which represent common combining ability profiles. A third cluster, Group 3, contained only the genotype P5, indicating its unique and significantly different combining ability pattern (Table 5; Fig. 3).

**Table 5:** Distribution of genotypes into different clusters based on HSGCA Value using K-Means Analysis.

| **Group 1** | **Group 2** | **Group 3** |
| --- | --- | --- |
| P1 P4 P10 P11 | P2 P3 P6 P7 P8 P9 | P5 |

**Fig 3:**
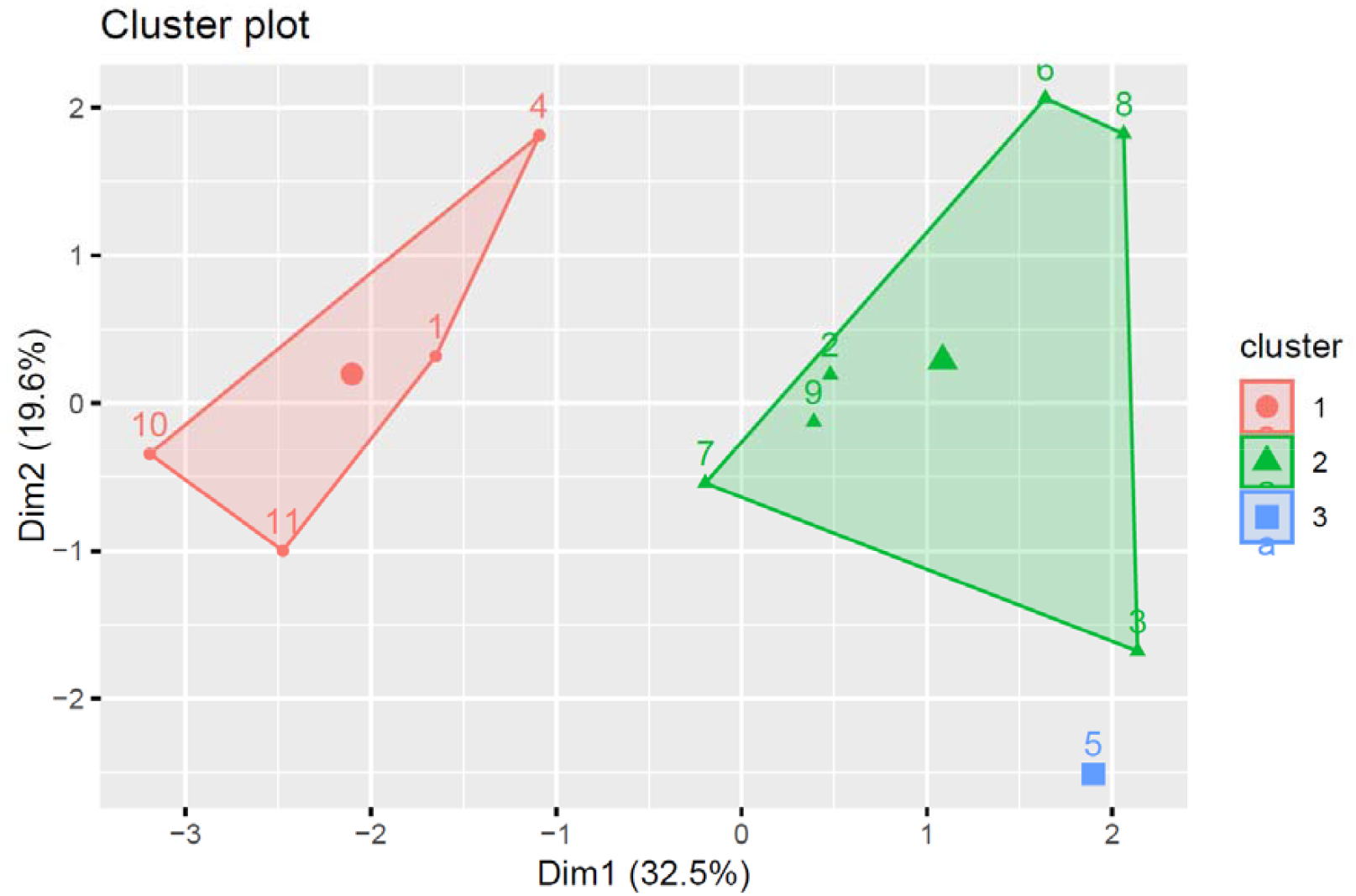
Grouping of 11 Genotypes into Three Clusters based on HSGCA (Combined GCA+SCA) using K-Means Analysis.

### 3.3. Validation of Heterotic Grouping Methods through Heterosis Analysis

The critical evaluation of the two clustering strategies—based on morphological traits versus combining ability (HSGCA)—was conducted by analyzing the heterosis expressed in crosses made within and between the derived groups. This validation aimed to determine which method more accurately predicts hybrid performance, with the fundamental hypothesis that a superior clustering approach would yield significantly higher heterosis in inter-group crosses compared to intra-group crosses. The predictive power of each method was rigorously assessed using both Mid-Parent Heterosis (MPH) and Better-Parent Heterosis (BPH).

#### Performance of Phenotypic Trait-Based Clusters

The heterosis analysis revealed significant inconsistencies in the predictive capability of the morphologically-derived clusters (Fig. 4 and Fig. 5). Contrary to theoretical expectations, crosses made within the same phenotypic cluster frequently exhibited substantial heterosis. For instance, the average MPH for intra-group crosses was remarkably high within Group 4 (29.6%) and Group 2 (22.42%) (Table 6). This paradoxical result indicates that genotypes grouped together based on phenotypic similarity were, in fact, genetically divergent for the loci governing yield and its component traits.

**Table 6:** Mid-Parent Heterosis (%) for crosses within and between phenotypic clusters.

| Mid-Parent Heterosis | Group 1 | Group 2 | Group 3 | Group 4 | Group 5 |
| --- | --- | --- | --- | --- | --- |
| **Group 1** | 4.64 | 4.12 | 17.6 | 14.04 | 16.5 |
| **Group 2** |  | 22.42 | 34.84 | 12.84 | 7.6 |
| **Group 3** |  |  | 0 | 49.5 | 15 |
| **Group 4** |  |  |  | 29.6 | 28 |
| **Group 5** |  |  |  |  | 0 |

**Fig 4:**
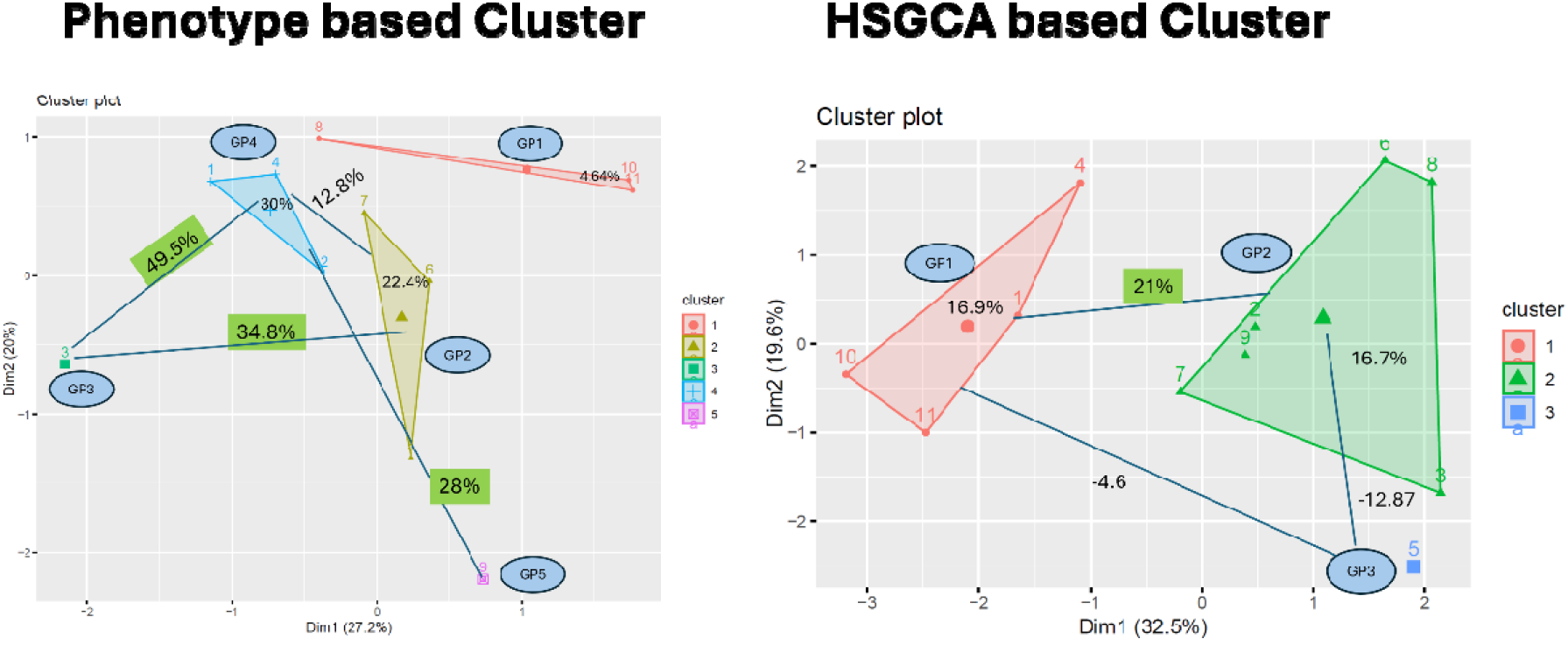
Validation of Phenotype-based vs. GCA-based Clustering Methods using Mid-Parent Heterosis.

**Fig 5:**
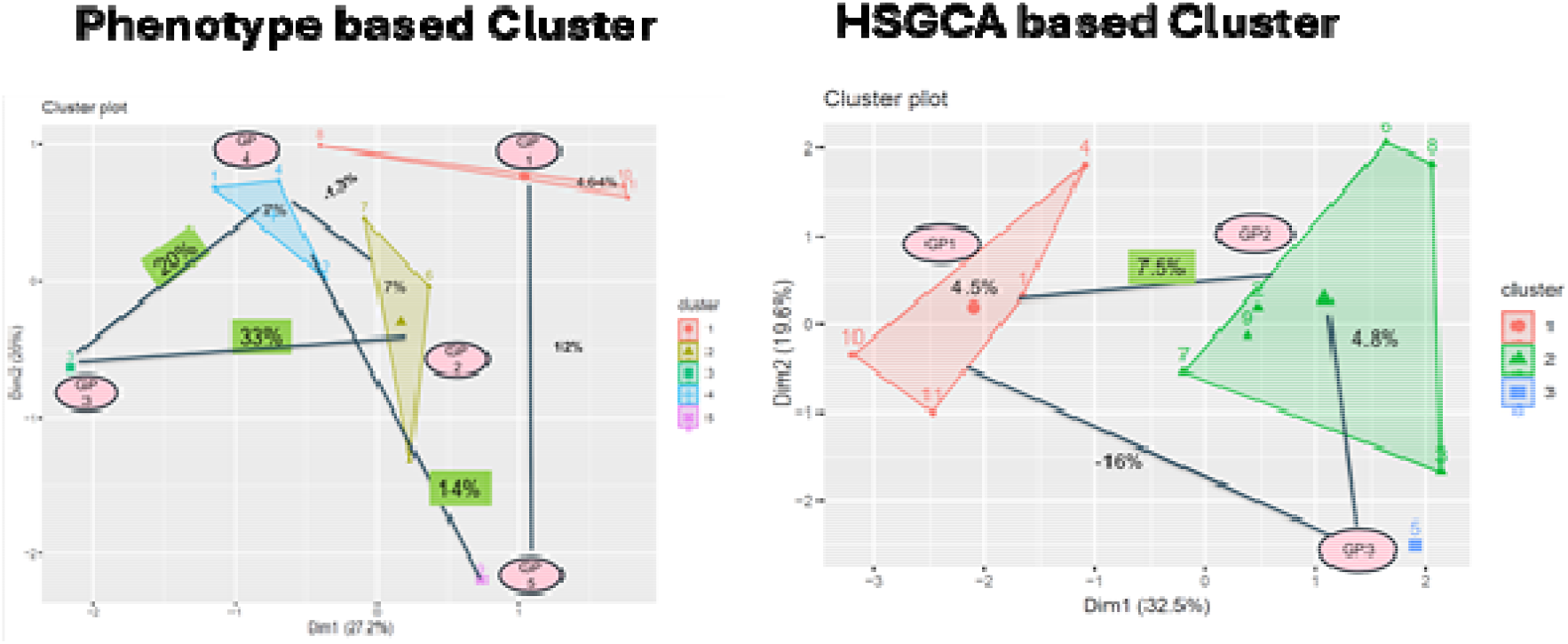
Validation of Phenotype-based vs. GCA-based Clustering Methods using Better-Parent Heterosis.

Furthermore, crosses between different phenotypic clusters failed to demonstrate a consistent heterotic advantage. The cross between Group 1 and Group 2 resulted in a very low MPH of 4.12%, a value significantly lower than the heterosis observed within either of those groups. While certain combinations, such as between Group 3 (P3) and Group 4, showed high heterosis (49.5% MPH), the overall pattern was erratic and unreliable for predictive breeding (Fig. 4). This inconsistency was further confirmed by the BPH analysis, where intra-cluster values were sometimes comparable to or even exceeded inter-cluster values (Table 7). These results demonstrate that clustering based on morphological data does not effectively capture the genetic divergence necessary for predicting heterosis.

**Table 7:** Better-Parent Heterosis (%) for crosses within and between phenotypic clusters.

| Better-Parent Heterosis | Group 1 | GROUP 2 | Group 3 | Group 4 | Group 5 |
| --- | --- | --- | --- | --- | --- |
| **Group 1** | 4.64 | -4.4 | 4.9 | 2.53 | 12.1 |
| **Group 2** |  | 6.9 | 20.02 | -1.3 | -1.5 |
| **Group 3** |  |  | 0 | 33 | 3 |
| **Group 4** |  |  |  | 6.8 | 14.05 |
| **Group 5** |  |  |  |  | 0 |

#### Superior Predictive Power of HSGCA-Based Clusters

In stark contrast, the heterotic groups defined by the HSGCA method exhibited a clear and logically consistent pattern (Fig. 4 and Fig. 5). The most significant finding was the pronounced superiority of inter-group crosses over intra-group crosses. The average MPH for hybrids between Group 1 and Group 2 was 20.89%, substantially exceeding the MPH observed within Group 1 (16.86%) and Group 2 (16.74%) (Table 8).

**Table 8:**
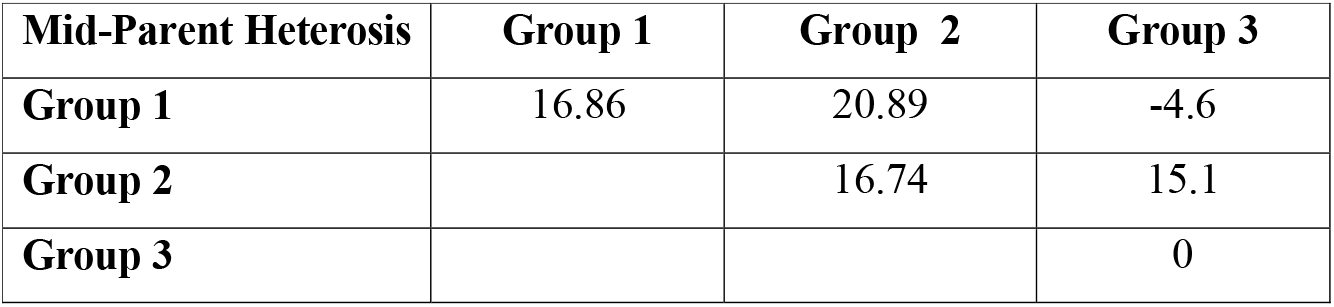
Mid-Parent Heterosis (%) for crosses within and between HSGCA-based clusters.

| **Mid-Parent Heterosis** | **Group 1** | **Group 2** | **Group 3** |
| --- | --- | --- | --- |
| **Group 1** | 16.86 | 20.89 | -4.6 |
| **Group 2** |  | 16.74 | 15.1 |
| **Group 3** |  |  | 0 |

This robust pattern was further validated by the more stringent BPH analysis. The BPH for inter-group crosses (Group 1 x Group 2) was 7.54%, significantly higher than the BPH within Group 1 (4.5%) and Group 2 (4.79%) (Table 9). The HSGCA method also provided a critical practical insight by clearly identifying genotype P5 (Group 3) as a poor combiner, as evidenced by the negative heterosis in its crosses with Group 1 (−4.6% MPH). This clear demarcation of heterotic patterns confirms that the HSGCA-based clustering successfully groups genotypes with similar combining ability, thereby creating genetically divergent pools between which crosses result in maximal heterosis.

**Table 9:** Better-Parent Heterosis (%) for crosses within and between HSGCA-based clusters.

| **Better Parent Heterosis** | **Group 1** | **Group 2** | **Group 3** |
| --- | --- | --- | --- |
| **Group 1** | 4.5 | 7.54 | -1602 |
| **Group 2** |  | 4.79 | 6.7 |
| **Group 3** |  |  | 0 |

## 4. Discussion

A comparative analysis of the two clustering methodologies revealed profound differences in their efficacy for predicting heterotic potential in bitter gourd. The initial approach, which relied on morphological trait-based clustering, proved to be an unreliable predictor of hybrid performance. This finding aligns with a substantial body of literature across various crops, where the correlation between phenotypic distance and heterosis has been consistently reported as weak and unpredictable (Dias et al., 2004; Yuan et al., 2008). A fundamental inconsistency in our study was the unexpectedly high level of heterosis observed within morphological clusters, a finding that directly contradicts the objective of creating genetically homologous groups. For instance, crosses within groups Group 4 and Group 2 yielded average mid-parent heterosis (MPH) values of 30% and 22.4%, respectively. This clearly indicates that genotypes exhibiting phenotypic similarity were, in fact, genetically divergent for critical yield-related loci, likely due to the influence of non-additive gene effects and epistasis that are not captured by morphology. Furthermore, the heterosis resulting from inter-cluster crosses was frequently inferior to that observed within single groups. The cross between Group 1 and Group 4, which produced an MPH of only 12.8%—significantly lower than the 30% heterosis within Group 4—exemplifies this failure. This inconsistent alignment with established heterotic principles confirms that phenotypic similarity is a poor indicator of the underlying genetic diversity necessary for hybrid vigor. An additional practical constraint of this method was the formation of two mono-genotypic clusters (Group 3 and Group 5). While these unique parents demonstrated high heterosis in specific combinations, their isolation is an unsustainable strategy for long-term population improvement, as it severely limits opportunities for genetic recombination.

In stark contrast, the heterotic grouping based on specific and general combining ability (HSGCA) method provided a vastly superior and more pragmatic framework. This finding strongly supports the growing consensus that breeding values are more direct predictors of hybrid performance than genetic distance measures (Fan et al., 2009; Adewale et al., 2023). By clustering genotypes according to their general and specific combining ability (GCA and SCA) values, this approach generated three robust groups with immediate practical utility. A key insight was the identification of parent P5 as a poor combiner, isolated in a singleton cluster (Group 3) that exhibited negative heterosis in crosses with other groups. This allows breeders to efficiently deprioritize such lines, thereby optimizing resource allocation. The most significant finding was the strong and consistent correlation between the genetic distance defined by combining ability and the resulting heterosis. The average MPH between the two primary clusters (Group 1 and Group 2) was 21%, substantially exceeding the heterosis observed within either group (16.9% and 16.7%, respectively). This clear discriminatory power was corroborated by the more stringent better-parent heterosis (BPH) analysis, where the inter-cluster value (7.5%) significantly outperformed the intra-cluster values. This demonstrates conclusively that the HSGCA method effectively partitions parental lines into genetically distinct heterotic pools, establishing inter-cluster crosses as a highly reliable strategy for maximizing hybrid performance. The success of this method in bitter gourd echoes its high accuracy and breeding efficiency reported in maize (Adewale et al., 2023) and baby corn (Kumar et al., 2022), suggesting its broad applicability.

## 5. Implications, Limitations, and Future Prospects

The superior performance of the HSGCA method can be attributed to its direct assessment of the additive (GCA) and non-additive (SCA) genetic variances that are the fundamental components of heterosis. Unlike morphological or molecular distance, which may include variation from neutral or non-contributing loci, combining ability values inherently reflect the actual gene action affecting the target traits, in this case, yield. However, it is important to acknowledge the limitations of this study. The HSGCA method requires the creation and evaluation of a large number of crosses (e.g., a diallel), which is resource-intensive. Future research could explore the integration of molecular markers with combining ability data to develop predictive models that might reduce the need for extensive field testing in subsequent breeding cycles. Furthermore, the stability of these heterotic groups across different environments should be investigated to ensure their robustness. Despite these considerations, our results provide a compelling evidence-based strategy for bitter gourd breeding. The clear delineation of heterotic groups (Group 1 and Group 2) offers a practical roadmap for breeders to systematically exploit heterosis, thereby accelerating the development of high-yielding hybrids and enhancing genetic gain in this important crop.

## 6. Conclusion

This study provides conclusive evidence that morphological diversity is an inadequate basis for predicting hybrid performance in bitter gourd, as it fails to capture the genetic determinants of heterosis. In contrast, the HSGCA method, which classifies genotypes based on their general and specific combining abilities, offers a scientifically robust framework for establishing meaningful heterotic groups. The efficacy of this approach is demonstrated by the clear heterotic pattern observed: crosses between the two primary groups (Group 1 and Group 2) consistently yielded significantly higher heterosis than crosses within the same group. This finding provides bitter gourd breeders with a validated, data-driven strategy for hybrid development. By strategically utilizing the defined heterotic pools, breeding programs can systematically enhance hybrid vigor, improve selection efficiency, and accelerate genetic gains in this important vegetable crop.

